# The mating ritual of a rainforest tinamou, *Tinamus major*

**DOI:** 10.1101/2024.04.05.588181

**Authors:** Yinger Guo, Jay J. Falk, Jorge L. Medina-Madrid, Silu Wang

## Abstract

The mating systems of many tropical birds remain poorly understood, especially those that elude observation in dense rainforest understories. Here we dissect mating behaviors between two Great Tinamous, *Tinamus major* (Tinamidae), a secretive species with a wide range across the lowland Central and South American tropics. Despite anecdotal preconceptions that males court the females, we observed courtship displays solely from the putative female tinamou. In this mating ritual recorded in their natural habitat, we observed clucking and soft-rolling songs along with a suite of courtship displays, such as neck-elongation, tail-raising, crouching, and feather-fluffing performed by the putative female. In contrast, the putative male watched while standing still, approached, walked away, and carried out a few mounting attempts. The clucking and soft-rolling songs sung by the putative female were of lower frequency, energy, and amplitude, but higher entropy than the common morning territorial songs recorded in the vicinity. The clucking song by the putative female was not previously described but we repeatedly observed this song type immediately before the soft-rolling songs during the behavioral displays. Clucking was of lower frequency and energy, and around ten times shorter than the soft-rolling songs. This behavioral auditory analysis enriches understanding of rainforest tinamou mating rituals and motivates scrutiny of the partition of courtship roles between sexes in a male parental-care mating system with little sexual dichromatism and reversed sexual size dimorphism.

## Introduction

There is limited understanding of the mating behavior of tinamous (Tinamidae) in their natural habitats (Brennan 2004; Schelsky 2004). *Tinamus major* is one of the most widely distributed tinamou species in Neotropical rainforests (Davies 2002; Li et al. 2023). The mating ritual of this species has only been described in captivity (Bonilla-Sanchez 2008), where mating behavior resembled the mating interaction of a congener, the Grey Tinamous (*T. tao*) (Solano-Ugalde et al. 2018). However, mating rituals in the wild may differ in complexity. Careful observation and analysis of a tinamou mating ritual in the wild would allow us to understand this elusive rainforest understory paleognath. Here we provide a detailed description and analysis of a recorded mating interaction between two *T. major* in Gamboa, Panamá.

## Methods

### Location

At 10:42 AM, on November 6, 2023, in Gamboa, Panama (9° 7’ 2.28” N, 79° 41’ 43.44” W), SW incidentally recorded the interaction of two *Tinamus major* individuals for 11 minutes and 12 seconds before an accidental interruption. We analyzed the recording to dissect the behavioral interaction. Because the birds were unmarked and sexual dimorphism in *T. major* is subtle, we could not visually determine the sex of the two individuals. Instead, we traced each individual throughout the recording, and classified putative sexes based on their distinctive positions during mounting in Tinamidae, where the male stands on top of the females (Beebe 1925; Bonilla-Sanchez 2008; Solano-Ugalde et al. 2018).

### Mating Behavior and Songs

To characterize the behavioral interaction during the mating ritual, we constructed an ethogram (**Table 1**) and provided examples in which each behavioral variable occurred during the recording.

**Table 1.** Ethogram of the mating ritual of *Tinamus major*. The example snapshots in **Fig. S1** in which the behavior occurred are listed in the last column.

| Putative sex | Category | Behavior | Description | Fig. | Time Occurred |
| --- | --- | --- | --- | --- | --- |
| Putative female<br>(yellow) | Vocal | *Soft-rolling song* | Soft-rolling song with syllables around 0.7s long, maximum amplitude ~ 490 db, and low frequency ~ 1153 Hz. | 4, 5 | Continuous |
|  | Vocal | *Clucking* | Short soft clucking-like notes, each lasts for around 0.7s, with maximum amplitude ~ 316 db, and low frequency ~ 1051 Hz. | 4, 5 | Continuous |
|  | Posture | *Tail-raising* | Raising the tail and flaring light undertail coverts while standing | 1, S1 | 00:28;01:13; 02:41-03:06; 03:26; 03:57; 08:24 |
|  | Posture | *Neck-stretching* | Enlongating the neck horizontally | 1, S1 | 02:33-02:41; 03:39; 08:53; 10:48 |
|  | Posture | *Crouching* | Lowering the body to the ground, like an egg-laying position | 1, S1 | 3:36-3:53; 04:01; 04:18; 10:32 |
|  | Posture | *Wing-drooping* | Lowering and spreading the primary feathers | 1 | 01:44-02:41 |
|  | Movement | *Feather-fluffing* | Fluffing up whole-body feathers, then gentling flapping the wings | 2 | 03:24-03:31 |
|  | Movement | *Following* | Moving towards the other bird, while the other was moving away | S1 | 06:49-06:58 |
| Putative male<br>(turquoise) | Movement | *Approach* | Moving towards the other bird, while the other was not moving away | 1 | 2:33 |
|  | Movement | *Flank-inspecting* | Within 0.5m, facing other bird's flank | 1, S1 | 3:24; 4:24; 10:48 |
|  | Movement | *Mouth-opening* | Leaning towards the other bird while opening its mouth | 1 | 1:44 |
|  | Movement | *Mount attempt* | Jumping over and briefly touching the other bird's back with feet | 3 | 3:57; 04:29 |
|  | Movement | *Standing atop* | Standing on the back of the other bird | 3 | 7:14 |

To understand song variations during the mating ritual, we analyzed all the syllables during 0:42-2:32 (**Fig. S1**) of the recording when the tinamous were close to the recorder so that the sound quality was sufficient for song analysis. We quantified entropy, energy, delta time (duration of the syllable), minimum and maximum entropy (the amount of disorder in the spectrum), maximum amplitude, sound exposure level (SEL), as well as minimum, peak, and maximum frequency for each syllable with RavenPro (Charif et al. 2010). To best capture the structural variation among syllables, we also derived a song variable called Relative Peak Position to identify the time at which the peak frequency occurred relative to the total duration of each syllable as (Peak time -start time)/(end time – start time).

To test whether there was a difference in the quantitative features of clucking versus soft-rolling songs, Linear Mixed Effect Models were used to examine the difference between the two song types with the *lmer* function in the *lme4* package (Bates et al. 2015) in R (R Core Team 2023). In each model, the response variables were quantitative measures of songs (e.g. frequency, energy, or entropy), the fixed effect was song types (clucking versus soft-rolling), and the random effect was song bouts (**Fig. 5**). We compared the full models to the corresponding null models with the same response variable and random effect, but no fixed effect.

We further compared clucking and soft-rolling mating syllables to regular songs recorded during the morning chorus in the vicinity. We downloaded the nearest (3.3 Km away) recording XC363937 (9.1332° N, -79.7195° W) from the incidental observation from Xeno-Canto (www.xeno-canto.org). This recording was recorded at 6:04 AM on April 7^th^, 2017, and lasted for 95.117 seconds containing 58 syllables. We compared the quantitative features among the three song types (clucking, soft-rolling, and regular) with principal component analysis. To understand specific variation among the three song types, we used linear mixed effect models with quantitative syllable measurements (e.g. frequency, energy, entropy) being response variables, the fixed effects being song types, and random effects being different sing bouts from different individuals.

## Results

The tinamous demonstrated several behavioral characters that were previously described in other tinamou species, such as *tail-raising* (**Table 1, Fig. 1**), *neck-stretching* (**Fig. 1**), *crouching* (**Fig. 1**), *wing-drooping* (**Fig. 1**), *following* (**Fig. S1**), and *soft-rolling songs* (**Table 1, Fig. 4-6**) by the putative female; as well as *approaching*, and *flank-inspecting, mount attempts, standing-atop* from the putative male. Although we did not observe copulation in this recording, we described two additional mating displays by the putative female: *feather-fluffing* (**Table 1, Fig. 2**) and *clucking* (**Table 1, Fig. 4-6**).

**Fig. 1.**
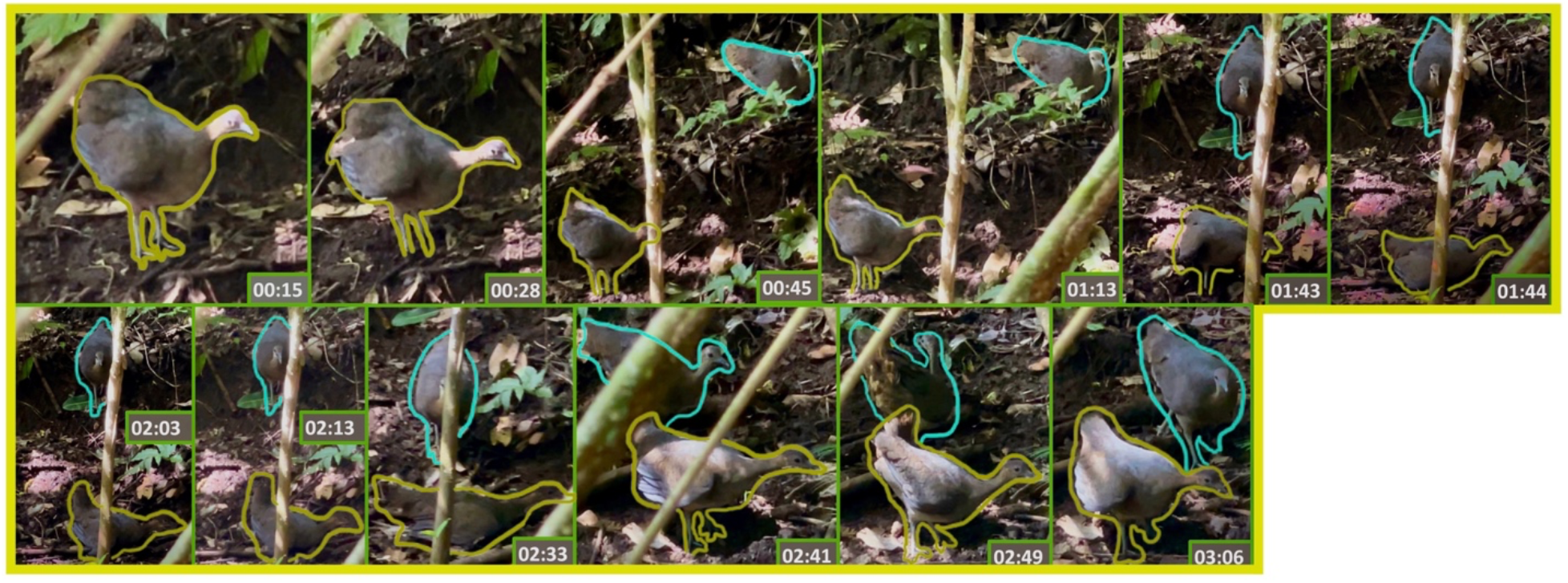
Putative female displays: neck-stretching, tail-raising, and crouching (**Table1**).

**Fig. 2.**
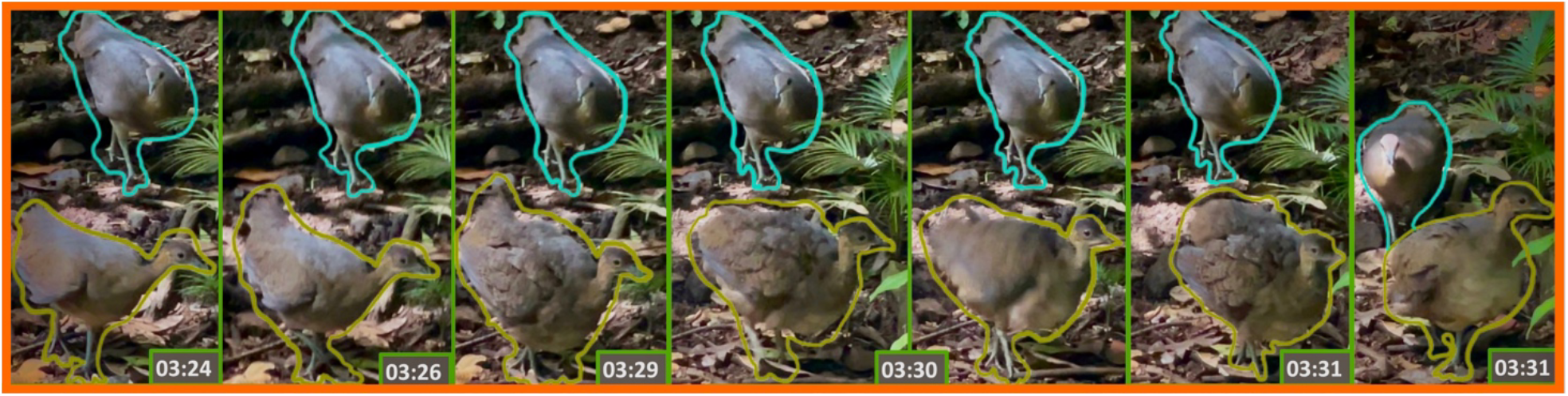
The putative female display: feather-fluffing (Table 1). The putative female drastically enlarged her apparent body size during this behavior. No preening was observed before or after this behavior. Feather-fluffing is immediately followed by a male approach and subsequent mounting attempt (**Fig. 3**).

**Fig. 3.**
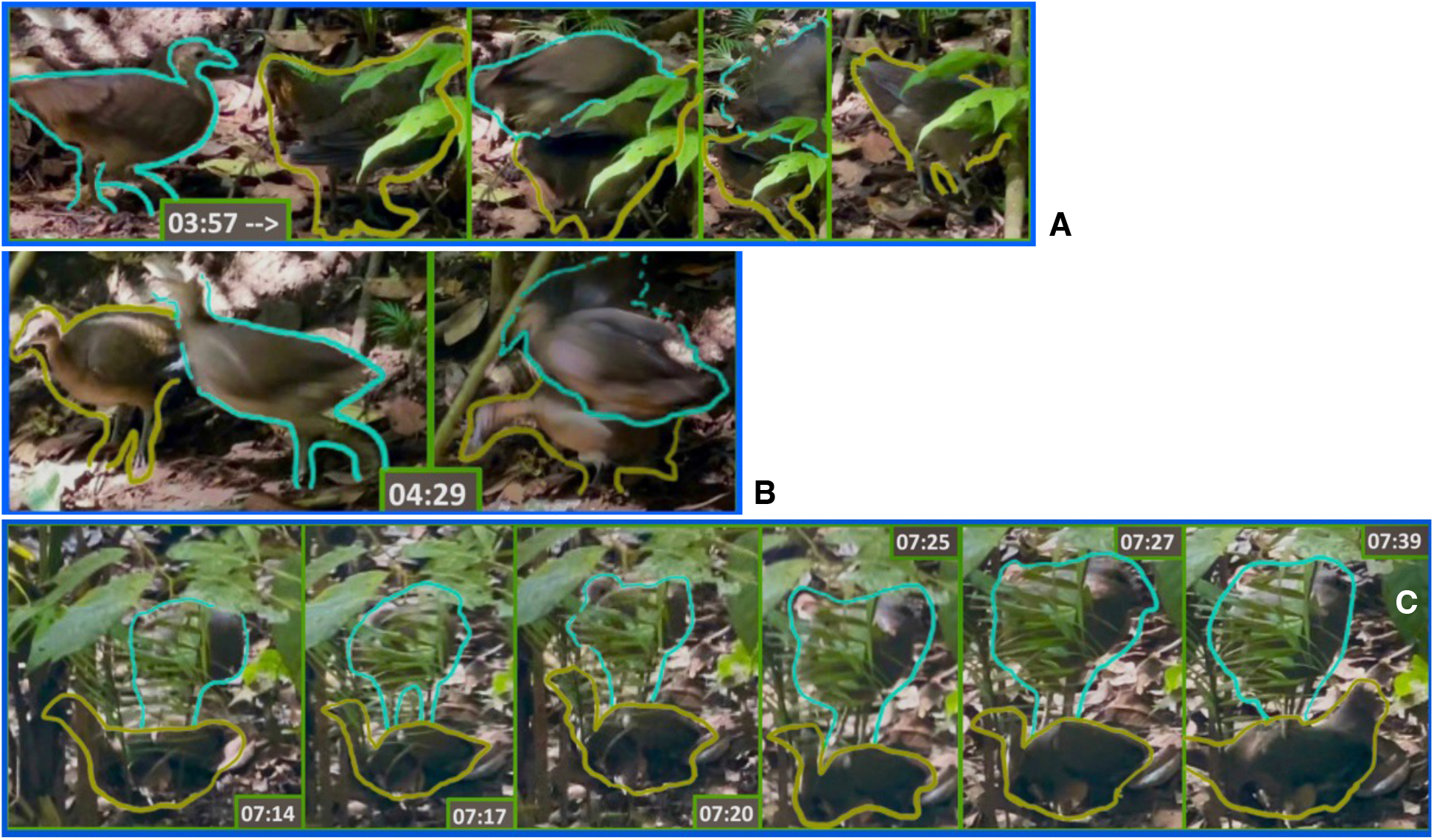
Putative male mounting attempt. The three times in which the putative male exhibited mounting attempts (**Fig. S1**).

**Fig. 4.**
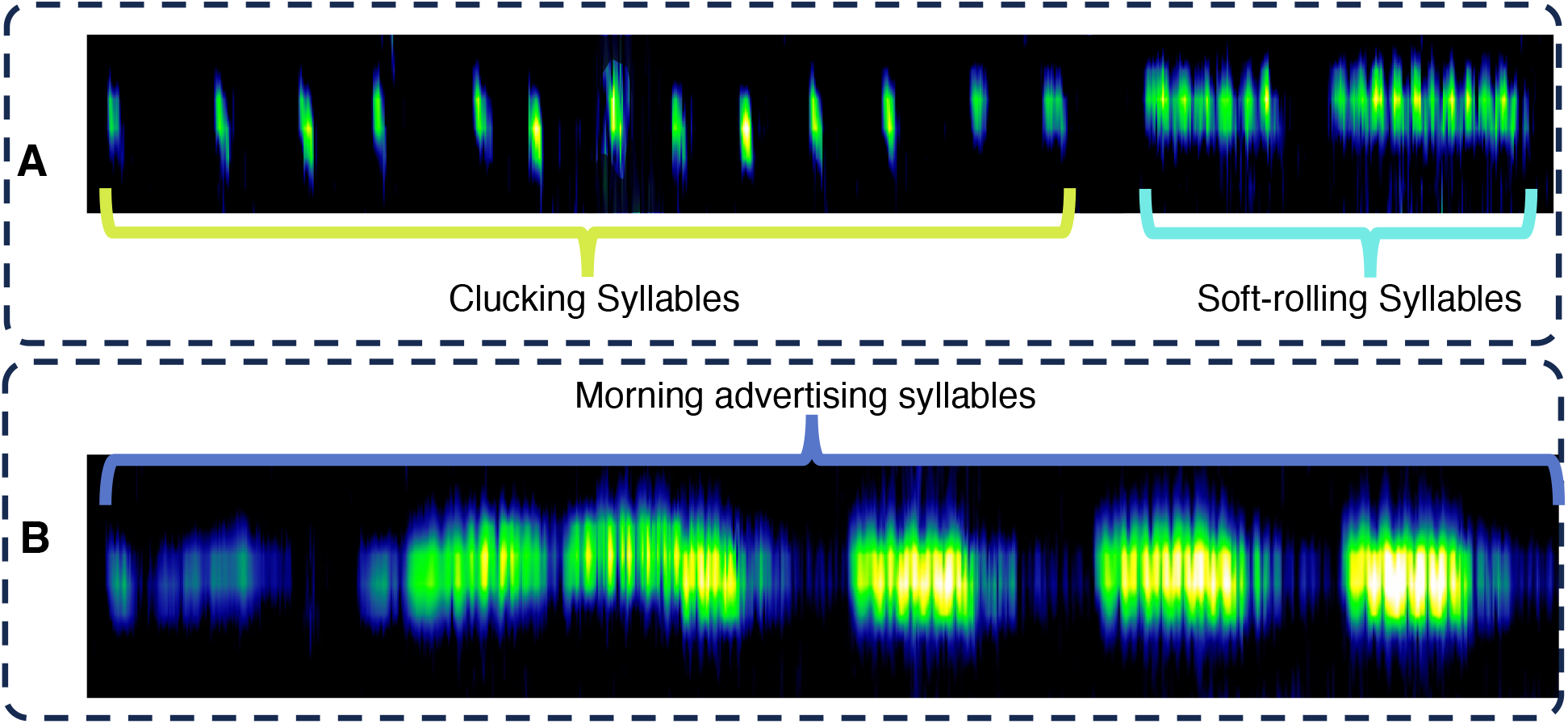
Spectrograms of clucking (yellow), soft-rolling songs (turquoise), and regular songs (dark blue). **A**, example spectrograms from song recording during the incidental observation of *T. major* mating ritual. **B**, example spectrograms of regular morning advertising songs.

**Fig. 5.**
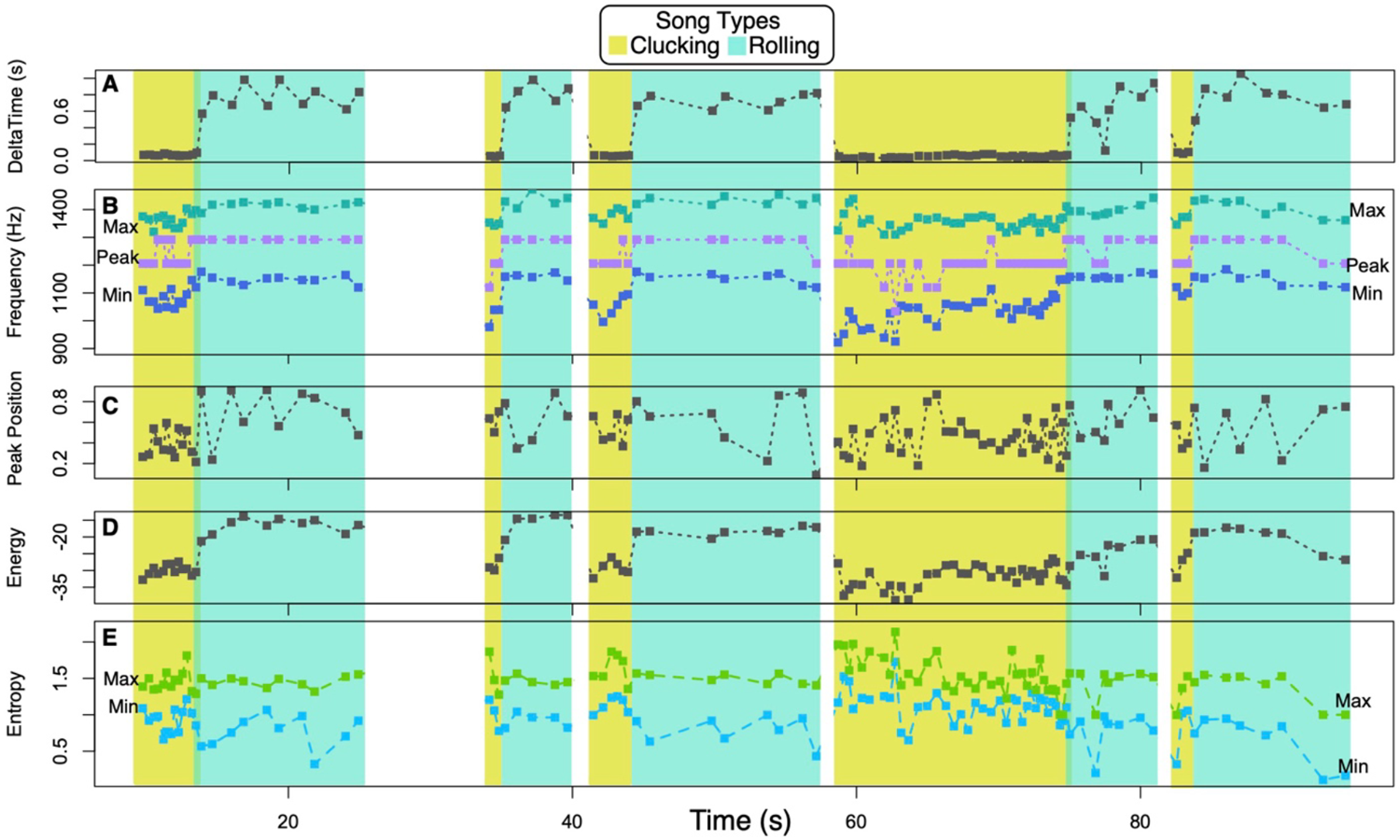
Quantitative features of putative female vocalization during mating ritual. There was significantly lower delta time (**A**), frequency (**B**), relative peak position (**C**), and energy (**D**) as well as higher entropy (**E**) in clucking (yellow) versus soft-rolling songs (turquoise). All the songs in the duration of 110 seconds are quantified here.

There was a significant difference between clucking and soft-rolling songs. The clucking songs showed lower delta time, minimum and maximum frequency, energy, peak Position, and greater minimum entropy (*p* < 0.001; **Fig. 5-6**). There was no significant difference in maximum entropy (*p* > 0.05; **Fig. 5-6**). The soft-rolling syllables were intermediate (**Fig. 6 A**) between morning advertising syllables and clucking syllables in serval audio measurements: high frequency (**Fig. 6 E**), energy (**Fig. 6 F**), and SEL (**Fig. 6 J**). The soft-rolling songs were similar to morning advertising songs in delta time (**Fig. 6 B**), peak (**Fig. 6 C**), and low frequency (**Fig. 6 D**). Soft-rolling songs are intermediate between clucking and morning advertising songs (**Fig. 6 A**). The two mating syllable types (clucking and soft-rolling) are both lower in Entropy (**Fig. 6 G, H**) and amplitude (**Fig. 6 K**) than the morning advertising syllables. Clucking and morning advertising songs both had relative peak positions around the center of the syllables while soft-rolling syllables tended to peak towards the end of the syllables (**Fig. 6, I**).

**Fig. 6.**
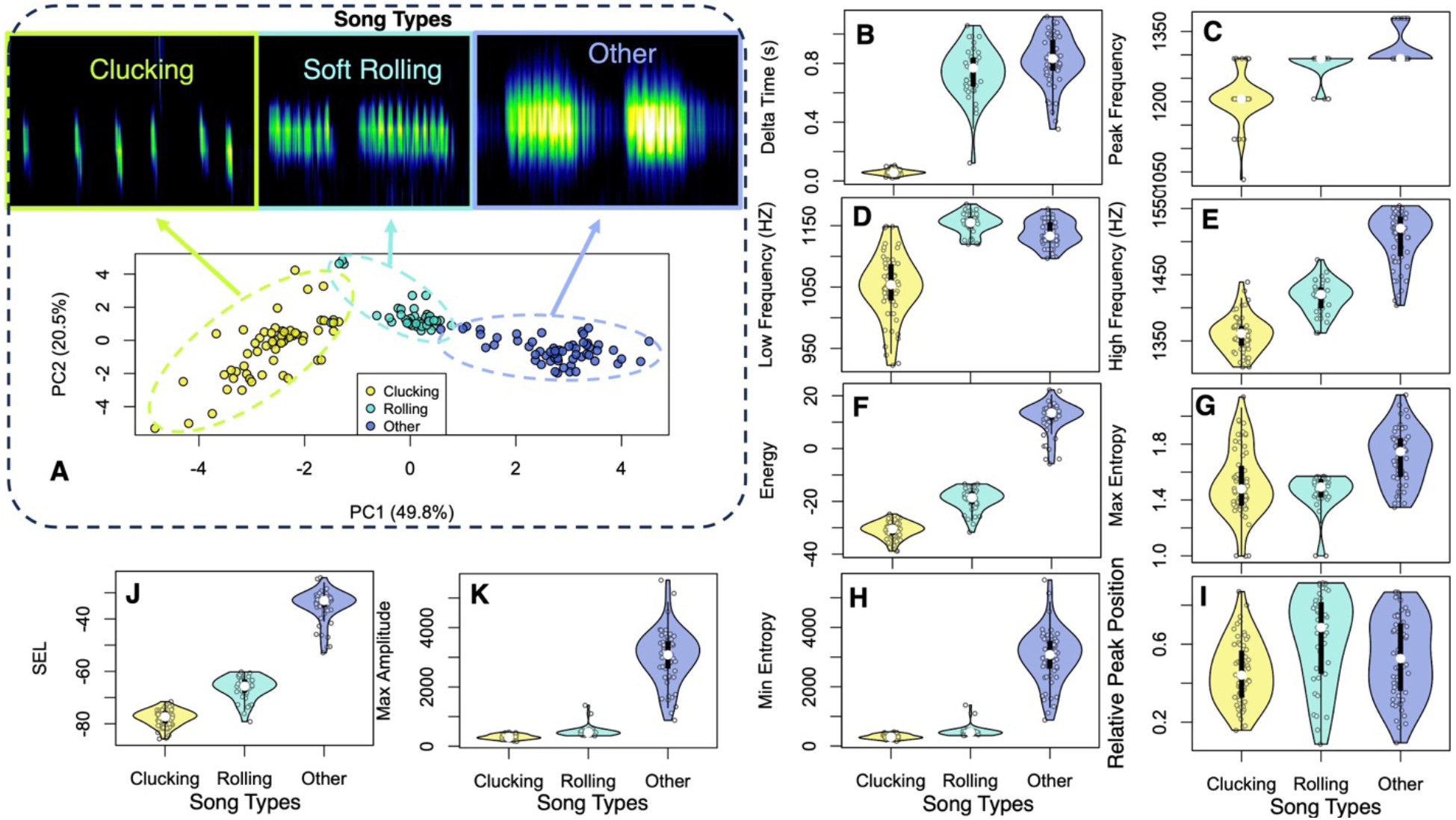
Auditory comparison among song types. The clucking and soft-rolling songs from the putative female is different from the morning advertising songs (other) recorded in the vicinity (3 Km apart). **A**, principal component analysis of song variables among the three song types. **B**-**I**: Violin plots of song variables among song types.

## Discussion

This incidental observation provides one of the first detailed descriptions of the *Tinamus major* mating ritual in their natural habitat. The behavioral interaction was similar to the mating ritual observed in captive *T. major* (Bonilla-Sanchez 2008) and wild *T. tao* (Solano-Ugalde et al. 2018), where the putative female demonstrated neck-stretching, and crouching, while the putative male stood on the back of the putative female.

We observed several display behaviors (**Table1, Fig. 1-3**) by the putative female (**Table 1, Fig. 1, 2**). *Similar to Tinamus tao*, we observed soft-rolling songs (**Fig. 4-6**), neck-stretching (**Fig. 1, S1**), crouching(**Fig. 1, S1**), wing-drooping(**Fig. 1, S1**), and tail-raising (**Fig. 1, S1**) by the putative female. The putative male did not show any display-like behavior (**Fig. S1**), was mostly watching while standing still (**Fig. 1-2**), then approached (Fig. 2), did flank-inspecting (**Fig. 1, S1**), mounting attempts (**Fig. 3, S1**), and standing atop (**Fig. 3, S1**) (**Table 1**). Female tail-raising was also observed in *Crypturellus variegatus* (Beebe 1925).

We also observed a previously undescribed behavior in the putative female, feather-fluffing (**Table 1, Fig. 2**). This form of display drastically enlarged the apparent size of the putative female, which exaggerated the sexual size dimorphism in *T. major* (Davies 2002). Feather-fluffing might occur together with neck-stretching (**Fig. 1**) which also enlarges apparent female body size. Female body size enlargement could be attractive to males, as immediately after the feather-fluffing, the putative male approached (**Fig. 2**) and then performed the first mounting attempt (**Fig. 3**), despite the prolonged hesitation before the feather-fluffing display (**Fig. S1**). More future observations in *T. major* and other tinamou species are needed to evaluate the prevalence of this type of display.

The putative female was the sole singer throughout the mating ritual with a complex suite of display-like behaviors (**Table 1, Fig. 1-2, S1**). The clucking and soft-rolling songs during mating were distinct from the morning advertising songs with lower frequency, energy, amplitude, and entropy (**Fig. 6**). The morning advertising songs are more commonly heard and are distinctive from other sympatric tinamou species (Bertelli and Tubaro 2002). To our knowledge, this is the first observation of the clucking syllables in the tinamou mating ritual. This type of syllable occurred quite frequently (∼34 times/minute) and occurred immediately before the soft-rolling songs (**Fig. 4-6**). Clucking is much shorter, with lower frequency and energy but higher entropy than soft-rolling songs (**Fig. 4-5**) during sexual displays (**Fig. 1-2**), which might prime the motivation state of sexual interactions. The soft-rolling songs resemble the regular advertising songs (**Fig. 4, 6**), but exhibited lower energy, entropy, and frequency. The intervals among clucking were uneven, and the counts of clucking before soft-rolling songs were variable (**Fig. 4, 5**).

The clucking, soft-rolling, and advertising songs that we compared (**Fig. 4, 6**) might represent female versus male song dimorphism of *T. major*. However, *T. major* females might also sing the morning advertising song type. Although there is limited understanding of song sexual dimorphism in *T. major*, song sexual dimorphism has been postulated in various species from its closely-related genus, *Crypturellus* (Boesman et al. 2018). In *C. brevirostris, C. cinereus*, and *C. soui*, the primary versus secondary songs were thought to be sung by territorial males versus paired females during mating duets. Similar to our observation, the primary songs, or morning advertising songs, were of higher frequency, more species-specific, and more frequently heard than the secondary songs (Boesman et al. 2018), or clucking and soft-rolling songs in this study. However, no video recording was paired with the putative duetting song analysis by Boesman et al. (2018), thus ethotypic sexing was impossible. Multimodal observations have been scarce because of the secretive behavior of the rainforest tinamous, which constrains understanding of the song dimorphism of rainforest tinamous.

In contrast to the preconception that males court the females in *Tinamus* (Del Hoyo et al. 2013), here we found that the putative female predominantly courted with a complex suite of displays, while the putative male seemed passive (**Fig. 1-3, S1**). However, the Tinamus male-display preconception was mostly based on anecdotes (Davies, 2002), which could have miscategorized the sexes. Because mating behavior is poorly understood in rainforest tinamous (Brennan 2004; Schelsky 2004) and there is a lack of obvious sexual dimorphism, we should be cautious with sexing or even putative sexing based on ethotypic assignment. Our observation of female-only courtship agreed with the rare field observations of mating rituals in *T. tao* (Solano-Ugalde et al. 2018) and *C. variegatus* (Beebe 1925), which collectively raised scrutiny in the allocation of courtship labor between sexes. The female premating efforts could be selected when male post-mating reproductive investment overweighs the total female reproductive investment (Andersson and Iwasa 1996). The *T. major* mating system in which males are the primary parental-caregiving sex could select for female courtship and male choosiness, aligning with the reversed sexual size dimorphism in this taxon.

## Acknowledgment

We thank Dahong Chen and members of the Forest Speciation Lab for helpful discussions, the Panama Ministry of Environment and Smithsonian Tropical Research Institute for the study permit, IACUC of the State University of New York at Buffalo, and the University of Colorado, Boulder for animal care guidelines. Funding was provided by SUNY Research Foundation to SW.

---

**Fig. S1.**
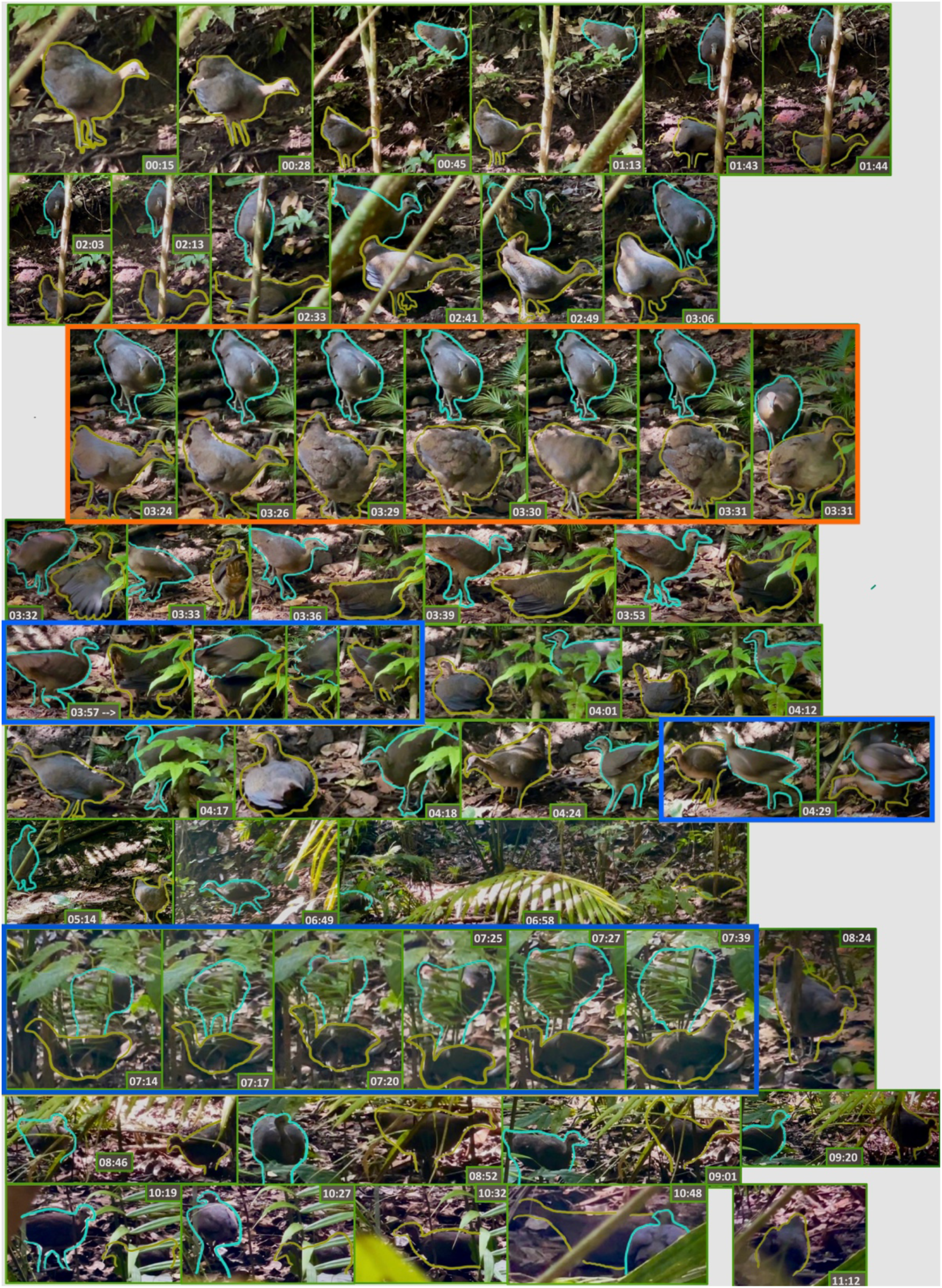
The mating ritual of *Tinamus major*. The putative female is contoured in yellow; the putative male is contoured in cyan.

